# Mapping Protein-Protein Interactions Using Data-Dependent Acquisition Without Dynamic Exclusion

**DOI:** 10.1101/2022.02.15.480563

**Authors:** Shen Zhang, Brett Larsen, Karen Colwill, Cassandra J. Wong, Ji-Young Youn, Anne-Claude Gingras

**Author notes:** Corresponding Author. Address: Lunenfeld-Tanenbaum Research Institute, Mount Sinai Hospital, 600 University Avenue, Toronto, ON, M5G 1X5, Canada.

## Abstract

Systematic analysis of affinity-purified samples by liquid chromatography coupled to mass spectrometry (LC-MS) requires high coverage, reproducibility, and sensitivity. Data-independent acquisition (DIA) approaches improve the reproducibility of protein-protein interaction detection by alleviating the stochasticity of data-dependent acquisition (DDA). However, the need for library generation and lack of multiplexing capabilities reduces their throughput, and analysis pipelines are still being optimized. In previous work using cell lysates, a fast MS/MS acquisition method with no dynamic exclusion (noDE) provided a comparable number of identifications and more accurate MS/MS intensity-based quantification than an optimized DDA method with dynamic exclusion (DE). Here, we have further optimized the noDE strategy for the analysis of protein-protein interactions and show that it provides better sensitivity and identifies more high confident interactors than the optimized DDA with DE and DIA approaches.

**TOC:** 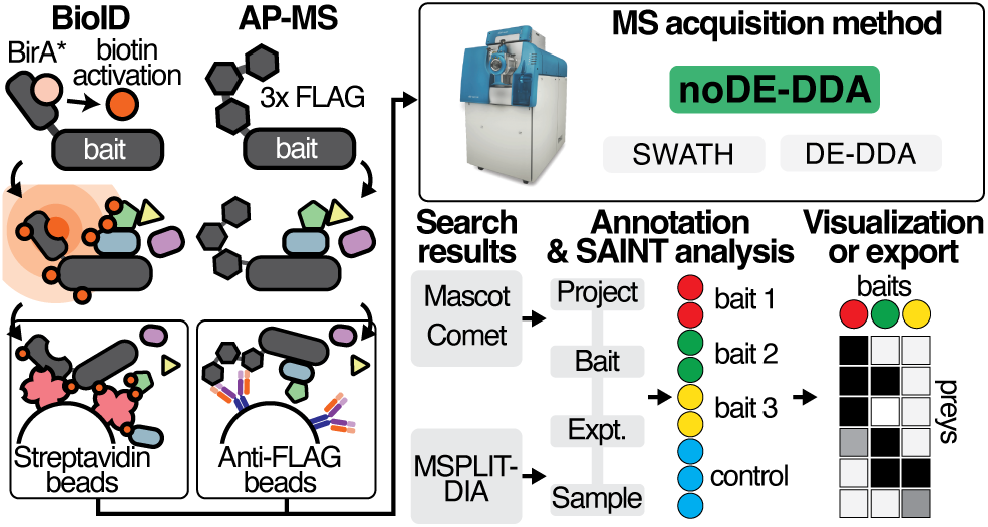

## Introduction

Protein-protein interactions are essential for all cellular functions and their contributions to disease phenotypes are being increasingly recognized^1, 2^. Protein interactions for a given protein of interest (bait) are typically identified by affinity purification (AP) of the bait and its stably associated binding partners^3^, or alternatively, by proximity-dependent biotinylation (e.g. BioID) of vicinal interactors^4^, that are subsequent captured on streptavidin supports^5,6^. Mass spectrometry (MS) based identification approaches are coupled to these purification methods. In the most extensively used MS strategy, known as shotgun or discovery proteomics, the MS instrument is operated in data-dependent acquisition (DDA), most often with dynamic exclusion (DE), which excludes re-selection of a peptide within a given time window^7^. Although this strategy improves the detection of low-level peptides in complex mixtures, the stochastic nature of precursor ion selection and under-sampling leads to a lack of reproducibility between samples^8^. We previously reported a method in which the DE step was omitted, resulting in improved quantification and reproducibility in isobaric tagging experiments with samples of moderate complexity^9^, but this method has not been applied to protein-protein interactions.

Alternatively, to alleviate the limitations associated with DDA with DE, strategies based on unbiased “data-independent acquisition” (DIA), in which every peptide within a specified window is fragmented, have been adopted^10, 11^. DIA can quantify the same number of proteins as typically identified by DDA methods, with better accuracy and reproducibility across many samples. However, there are limitations to DIA strategies. First, most require a spectral library—created via DDA analysis—for peptide quantification. Separating identification from quantification not only reduces the throughput, but also increases the possibility of mismatching between fragment ions in the DDA and DIA spectra. Although there are many publicly available libraries and retention time calibration strategies to address these issues^12-15^, studies have reported decreased identifications when using public libraries instead of sample-specific libraries^16-18^. Second, DIA spectra are complex, making accurate data extraction and analysis challenging, especially for some post-translational modifications^19^. Finally, an inherent problem caused by the wide isolation windows used in DIA experiments is their general incompatibility with isobaric labelling strategies, which require peptides to be individually isolated for accurate label quantitation.

In this study, we further optimized our fast MS/MS acquisition without DE (noDE-DDA) for AP-MS and BioID analyses and compared it to DDA with DE (DE-DDA) and the DIA approach SWATH (Sequential Window Acquisition of all Theoretical Mass Spectra)-MS methods. The optimized noDE-DDA strategy achieved better sensitivity and identified more high confident interactors than DE-DDA and provided comparable or better results than SWATH-MS^20^, depending on sample complexity. Importantly, this strategy can be implemented using standard informatics tools for peptide identification and quantification, facilitating data analysis.

## Experimental

### Sample Preparation

For the preparation of AP-MS (N-terminally 3×FLAG-tagged eukaryotic translation initliation factor 4A2 (EIF4A2) and methylphosphate capping enzyme (MEPCE)) and BioID (two N-terminally BirA-FLAG-tagged activated forms (G12V, Q61H) of KRAS proto-oncogene, GTPase (KRAS)) samples, see the Supporting Information.

### MS acquisition on TripleTOF mass spectrometers operated in DDA and DIA (SWATH-MS) modes

FLAG-AP and BioID samples were directly loaded at 800 nL/min onto 15 cm 100 μm ID emitter tips packed in-house with 3.5 μm Reprosil C18 (Dr. Maische). The peptides were eluted from the column at 400 nL/min over a 90 min gradient generated by a 425 NanoLC (Eksigent, Redwood, CA, USA) and analyzed on a TripleTOF 6600 instrument (AB SCIEX, Concord, Ontario, Canada). The gradient started at 2% acetonitrile with 0.1% formic acid and increased to 35% acetonitrile over 90 min, followed by a 15 min wash at 80% acetonitrile and a 15 min equilibration at 2% acetonitrile, for a total of 120 min.

For *Escherichia coli* lysate analysis, 500 ng MassPREP *E. coli* digestion standard (Waters, Cat#186003196) was loaded per injection. The noDE-DDA method was optimized by testing several acquisition parameters, including the use of fixed or rolling collision energy (CE); low resolution Q1 or unit resolution Q1; high resolution MS/MS acquisition mode or high sensitivity MS/MS acquisition mode; the number of peptide ions selected for MS/MS fragmentation (top 75 or top 100); the precursor intensity threshold (100 cps or 300 cps); and the MS/MS accumulation time (30, 45, or 60 ms).

Each AP-MS and BioID sample was analyzed on a TripleTOF 6600 in DDA mode with and without DE and in DIA (SWATH-MS) mode. DDA with DE consisted of one 250 ms MS1 TOF survey scan from 400–1250 Da followed by 10 × 100 ms MS2 candidate ion scans from 100–1800 Da in high sensitivity mode. Only ions with a charge of 2+ to 5+ that exceeded a threshold of 300 cps were selected for MS2, and former precursors were excluded for 7 s after one occurrence. For DDA mode without DE, precursors that exceeded a threshold of 100 cps were selected for MS2 and 100 × 30 ms MS2 candidate ion scans from 100–1800 Da were acquired in high sensitivity mode, using the same acquisition parameters otherwise. For SWATH-MS on the TripleTOF 6600, acquisition consisted of 55 variable windows covering a mass range of 400–1250 a.m.u. (cycle time ∼3.5 s). The MS1 and MS2 accumulation times were 250 ms and 65 ms, respectively.

### MS data analysis, interaction data analysis and visualization

For the DDA and DIA MS data analysis, interaction data analysis and visualization by SAINTexpress and ProHits-viz, see the Supporting Information.

## Results and Discussion

### Method optimization

We first optimized noDE-DDA on a SCIEX Triple TOF 6600+ instrument using *E. coli* lysate, which has a similar complexity to typical affinity purified samples from human cells. We systematically evaluated the influences of different parameters (fragmentation collision energy, isolation window width, MS/MS resolution, precursor intensity threshold, MS/MS accumulation time, and MS/MS candidate number; **Table S1**) on identification, and compared the results to those from a standard DE-DDA method^21^. Compared to DE-DDA (**Table S1**), most of the noDE-DDA methods yielded increased numbers of PSMs, peptides, and proteins and all methods increased sequence coverage. Among the noDE-DDA methods, the low resolution Q1 (i.e., a wider Q1 isolation window) and high-resolution MS/MS acquisition mode provided fewest identifications. The wider isolation window likely increased the number of chimeric spectra, lowering the spectrum score and thus decreasing identification. Although high-resolution MS/MS acquisition mode improved the accuracy of fragment ions, it sacrificed sensitivity, as counts were lower for all fragment ions compared to high-sensitivity MS/MS acquisition mode. The remaining noDE-DDA methods had minimal differences in protein and peptide identifications or sequence coverage, but had higher MS/MS acquisition numbers and lower MS/MS accumulation times in their MS parameter settings, providing more PSMs, which benefits spectral count-based quantification. In the following analyses, we used the noDE method with a 100 cps precursor intensity threshold, the top 100 MS/MS candidates, and a 30 ms MS/MS accumulation time, which identified the most proteins, peptides, and PSMs, and compared it to our DE-DDA and DIA methods.

### AP-MS sample identification

To test whether the noDE-DDA method was suitable for AP samples, we selected two well-characterized baits, EIF4A2 and MEPCE. EIF4A2, an RNA helicase, is a component of the eIF4F complex that associates with the mRNA 5′ cap, playing a crucial role in translation initiation. MEPCE is a 7SK snRNA methylphosphate capping enzyme, which stably associates with the 7SK ribonucleoprotein complex to regulate the activity of the p-TEFb complex (cyclin dependent kinase 9/cyclin T). These two proteins and the negative control (green fluorescent protein, GFP) were fused to a 3×FLAG tag at their N-terminus^21^. The fusion proteins were stably expressed in HEK293 Flp-In T-REx cells, affinity purified, and digested, and peptides were analyzed using the noDE-DDA, DE-DDA, and SWATH-MS acquisition methods in triplicate. For EIF4A2, noDE-DDE identified more PSMs (20180 vs. 8031, *p*-value = 0.0014), peptides (3799 vs. 3246, *p*-value = 0.0237), and proteins (1046 vs. 782, *p*-value = 0.0080) than the DE-DDA method (**Fig. S1A-C**), and more peptides (3799 vs. 3135, *p*-value = 0.0250) and proteins (1046 vs. 739, *p*-value = 0.0054) but fewer PSMs (20180 vs. 22456) than SWATH-MS analysis using the MSPLIT-DIA spectral matching approach. For MEPCE, noDE-DDE identified more PSMs (15918 vs. 7230, *p*-value = 0.0024), comparable peptides (3317 vs. 3272), and more proteins (1060 vs. 841, *p*-value = 0.0013) than the DE-DDA method (**Fig. S1D-F**), and less PSMs (15918 vs. 22710, *p*-value = 0.0020), comparable peptides (3317 vs. 3207), and more proteins (1060 vs. 807, *p*-value = 0.0009) than SWATH-MS analysis using MSPLIT-DIA.

We used SAINTexpress to compare samples to negative controls and assign confidence scores for true interactors^22, 23^. At a 1% Bayesian (B) FDR threshold, noDE-DDA identified 118 interactors for EIF4A2, 70% more than DE-DDA (69 interactors) and 18% more than MSPLIT-DIA (100 interactors) (**Fig. 1A**). The noDE-DDA method captured ∼80% of the interaction partners identified by DE-DDA (**Fig. 1B)**; and those only detected by DE-DDA did not have any reported interactions with EIF4A2 in the BioGRID repository^24, 25^. Conversely, 33 of the 48 preys uniquely identified by the noDE-DDA method have functions or activities related to EIF4A2. Five have reported interactions with EIF4A2 in BioGRID (CLNS1A, TPM3, EIF3G, PRRC2C, and PRRC2B), and 28 have RNA-binding activity, including 11 with roles in mRNA translation. Thus, the noDE-DDA method identified functionally relevant interactors of EIF4A2. MSPLIT-DIA identified 33 unique interactors, most with annotated RNA-binding activity and roles in mRNA translation, suggesting the complementarity of the MSPLIT-DIA and noDE-DDA methods.

**Figure 1.**
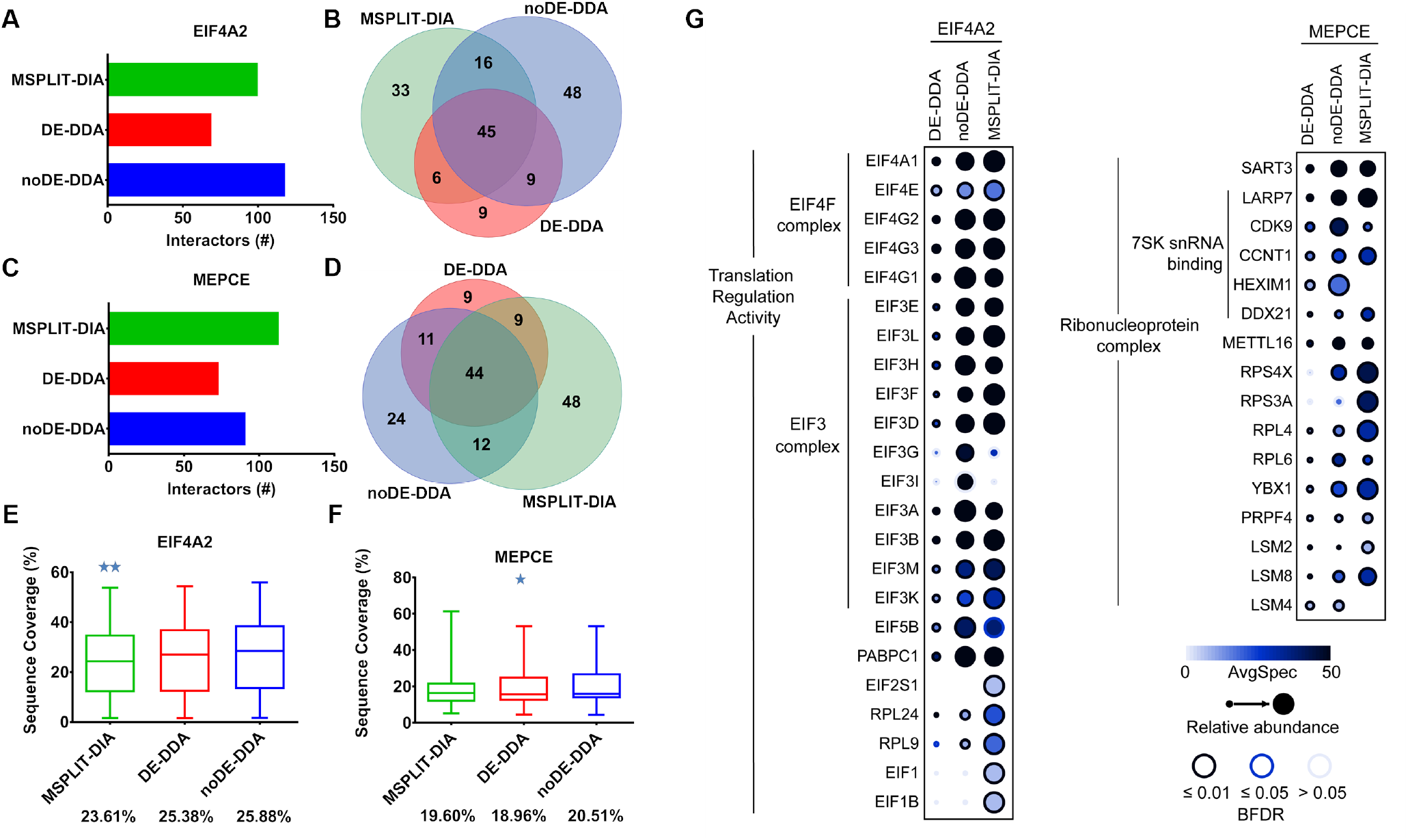
High-confidence AP-MS interactors of EIF4A2, MEPCE (A and C, respectively; SAINTexpress, BFDR ≤1%) and their overlap between different analysis methods (B and D). The sequence coverages of EIF4A2 and MEPCE interactors identified by all three methods are shown in E and F, respectively. \**p* < 0.05, \*\**p* < 0.01, by two-tailed paired Wilcoxon *t*-test. Spectral counts of a subset of known EIF4A2 and MEPCE signaling partners by AP-MS (SAINTexpress, BFDR ≤ 1%) are shown in G. The BFDR score is represented by the edge circle color; the color intensity indicates the average spectral count across replicates, and the size of the circle is proportional to the maximum value for a given prey across the entire dataset.

Similarly, SAINTexpress analysis of the MEPCE AP-MS noDE-DDA data identified 91 interactors, ∼25% more than DE-DDA (73 interactors) but ∼24% less than MSPLIT-DIA (113 interactors) (**Fig. 1C)**. Overlap analysis of the three methods showed that noDE-DDA captured ∼75% of the interaction partners identified by DE-DDA (**Fig. 1D**). In addition, 24 preys were uniquely identified by noDE-DDA, and 10 (∼40%) of these have reported interactions with MEPCE in BioGRID (CSNK2A1, HADHB, HNRNPA1, LRPAP1, RPS4X, TOP1, ALYREF, PRPF6, DDX5, SNRNP200), suggesting that the other preys identified by the noDE-DDA method are also true interactors of MEPCE.

We also investigated whether the noDE-DDA method could provide higher sequence coverage, which is required for the detailed characterization of specific protein regions, such as those harboring post-translational modifications (PTMs) and alternative splicing sites, which can increase our functional understanding of interaction networks. For EIF4A2 interactors identified by all three methods, noDE-DDA provided the highest average sequence coverage (25.88%), which was significantly higher than that of MSPLIT-DIA (23.61%, *p-*value *=*0.0013) (**Fig. 1E)**. For MEPCE, noDE-DDA also provided the highest average sequence coverage (20.51%), which was significantly higher than that of DE-DDA (18.96%, *p-*value *=*0.0311) (**Fig. 1F)**. In addition to increased identification and sequence coverage of interactors, the noDE-DDA method also had a higher range of PSM values than the DE-DDA method, with comparable range to the MSPLIT-DIA method (**Fig. S2**). The spectral counts of significant interactors identified by the different methods were significantly correlated (R^2^ > 0.895) (**Fig. S3**), and noDE-DDA yielded higher spectral counts per protein than DE-DDA (**Fig. S3A, B**) and comparable spectral counts to MSPLIT-DIA (**Fig. S3C, D**), and these trends were also detected for proteins functionally related to or known to stably associate with EIF4A2 or MEPCE (**Fig 1G**)^26^.

### BioID sample identification

Next, we tested if noDE-DDA improved peptide and protein identification with BioID samples, which have greater complexity than AP-MS samples. For this, we tested two activated forms of KRAS (G12V, Q61H) in an *in vivo* biotinylation experiment. For both KRAS G12V (**Figure S4A-C**) and KRAS Q61H (**Figure S4D-F**), noDE-DDA identified more PSMs, peptides, and proteins than DE-DDA and MSPLIT-DIA, suggesting that the advantages of noDE-DDA are more pronounced as samples increase in complexity.

When SAINTexpress (BFDR ≤ 1%) was applied to identify high-confidence proximal interactions for KRAS G12V, noDE-DDA identified 465 interactors, 35% more than DE-DDA (344 interactors) and 38% more than MSPLIT-DIA (337 interactors) (**Fig. 2A)**. The noDE method identified 95.35% and 91.99% of the interaction partners identified by DE-DDA and MSPLIT-DIA, respectively (**Fig. 2B)**. Of the 38 detected only by DE-DDA or MSPLIT-DIA, only 12 were reported KRAS interaction partners in BioGRID. In contrast, most of the 97 preys uniquely identified by the noDE-DDA method have functions or activities related to KRAS signaling, and 46 of them (∼47%) have reported interactions with KRAS in BioGRID.

**Figure 2.**
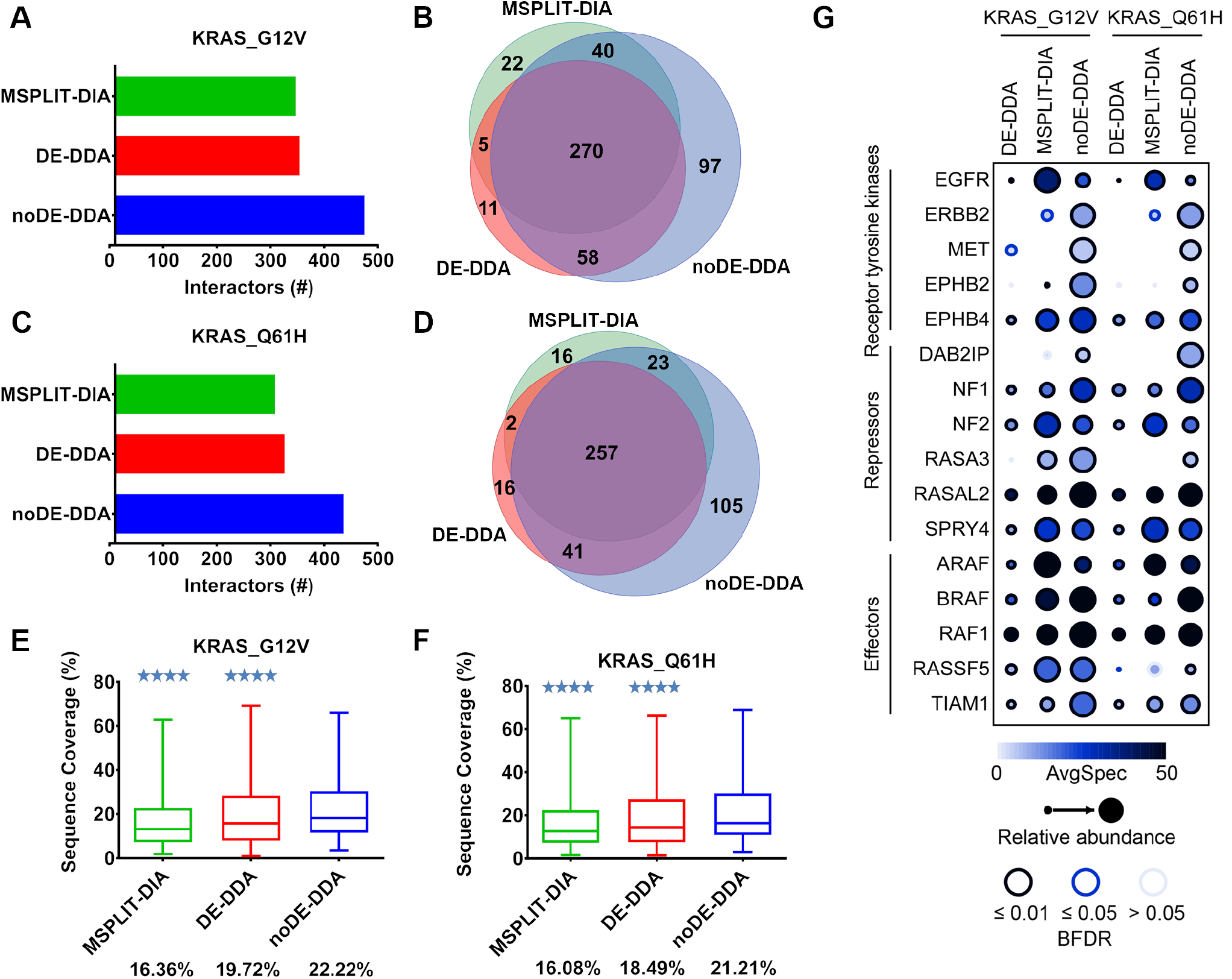
High-confidence BioID interactors of KRAS_G12V and KRAS_Q61H (A and C, respectively; SAINTexpress, BFDR ≤ 1%) and their overlap between different analysis methods (B and D). The sequence coverages of KRAS_G12V and KRAS_Q61H interactors identified by all three methods are shown in E and F, respectively. \**p* < 0.05, \*\**p* < 0.01, \*\*\**p* < 0.001, and \*\*\*\**p* < 0.0001, by two-tailed paired Wilcoxon *t*-test. Spectral counts of a subset of known signaling partners of KRAS G12V and Q61H by BioID (SAINTexpress, BFDR ≤1%) are shown in G. The BFDR score is represented by the edge circle color; the color intensity indicates the average spectral count across replicates, and the size of the circle is proportional to the maximum value for a given prey across the entire dataset.

Similarly, after SAINTexpress, noDE-DDA identified 426 interactors for KRAS Q61H, 35% more than DE-DDA (316 interactors) and 43% more than MSPLIT-DIA (298 interactors) (**Fig. 2C**). Comparison of the three methods showed that noDE-DDA detected 94.30% and 93.96% of the interaction partners identified by DE-DDA and MSPLIT-DIA, respectively (**Fig. 2D**). In addition, 105 preys were uniquely identified by noDE-DDA and 48 (∼46%) of these have reported interactions with KRAS in BioGRID. High-confidence noDE-DDA preys were enriched for gene ontology (GO) molecular function (MF) term “Ras GTPase binding” (G12V = 6^th^ rank, *p*-value 1.75E-25; Q61H = 7^th^ rank, *p*-value 1.10E-15), while these terms were enriched, but with lower ranks and p-vales in the DE-DDA (13^th^ rank, *p*-value 2.45E-07; 15^th^ rank, *p*-value 1.34E-07) and the MSPLIT-DIA (15^th^ rank, *p*-value 3.71E-08; 30^th^ rank, *p*-value 1.31E-05) for G12V and Q61H, respectively (**Table S2**)^27^. Together, this suggests that the noDE-DDA method identified more functionally relevant KRAS variant interactors than the other two methods.

Consistent with these findings, noDE-DDA provided the highest average sequence coverage for KRAS G12V and Q61H proximity interactors (22.22% and 21.21%, respectively), significantly higher than DE-DDA (19.72% and 18.49%, *p*-value<0.0001) and MSPLIT-DIA (16.36% and 16.08%, *p*-value<0.0001) (**Fig. 2E-F**) and this was associated with a higher range of PSM values than DE-DDA and comparable PSM range to MSPLIT-DIA (**Fig. S5**). For significant interactors identified by different methods, noDE-DDA provided much higher spectral counts than DE-DDA (**Fig. S6A, B**) and comparable spectral counts to MSPLIT-DIA (**Fig. S6C, D**). This is also evident by focusing on proteins known to be involved in RAS signaling (**Fig. 2G**), where noDE-DDA generally produced increased spectral counts for identified preys, and in some cases—such as DAB2 interacting protein, which activates RAS GTPases—noDE-DDA was the only method to identify the associated protein.

Together, these results show that for KRAS, the noDE-DDA method was superior to the other two methods, and suggest that when analyzing more complex enriched samples, noDE-DDA can identify more unique high-confidence interactors, with higher sequence coverage than the DE-DDA and MSPLIT-DIA methods.

## Conclusion

Here, we report the optimization and application of an efficient noDE-DDA method for quantitative interactome analysis using spectral counting. With all AP-MS and BioID samples tested, noDE-DDA identified more proteins and spectral counts per protein than a standard DE-DDA method, and recapitulated nearly all high-confidence interactors identified by DE-DDA. Compared with MSPLIT-DIA, the noDE-DDA method provided comparable and complementary results for less complex AP-MS samples. For more complex BioID samples, noDE-DDE identified almost all interactors identified by MSPLIT-DIA, as well as many unique true interactors. Furthermore, unlike DIA methods, noDE-DDA uses standard informatics tools for identification and quantification, and more importantly, this method is compatible with multiplex quantification, which is ideal for simultaneous sample measurements in dynamic interactome analyses.

## Associated Content

## Supporting information

The Supporting Information is available free of charge on the ACS Publications website at http://pubs.acs.org.

Details of sample preparation, MS detection, data analysis and visualization, identification of *E. coli* proteins with different MS parameters, GO enrichment analysis of KRAS_G12V and KRAS_Q61H interactors, the number of PSMs, peptides, and proteins identified in EIF4A2 and MEPCE AP-MS samples analyzed with the different methods, spectral count distributions of all high-confidence interactors for EIF4A2 and MEPCE, correlations between the log2-transformed spectral counts of all high-confidence interactors identified for EIF4A2 and MEPCE, the number of PSMs, peptides, and proteins identified in BioID samples of mutated KRAS after analysis by the different methods, spectral count distributions of all high-confidence interactors for KRAS G12V and KRAS Q61H, correlations between the log2-transformed spectral counts of all high-confidence interactors identified for KRAS G12V and KRAS Q61H.

## Notes

The authors declare no competing financial interest.

## Acknowledgments

We thank Dushyandi Rajendran for help preparing AP-MS samples, Kento T. Abe for help preparing TOC figure, and High-Fidelity Science Communications for editing the manuscript. S.Z. was supported by a Mitacs Elevate postdoctoral fellowship and an ASMS Postdoctoral Career Development Award. A.-C.G. is supported by a Canadian Institutes of Health Research Foundation Grant (FDN 143301) and an Ontario Research Fund Grant (ORF-RE08-065). Proteomics work was performed at the Network Biology Collaborative Centre at the Lunenfeld-Tanenbaum Research Institute, a facility supported by Canada Foundation for Innovation funding, the Ontarian Government, and Genome Canada and Ontario Genomics (OGI-139). A.-C.G. is the Canada Research Chair in Functional Proteomics and the Lea Reichmann Chair in Cancer Proteomics.

