## Supplementary documents for "Mapping Protein-Protein Interactions Using Data-Dependent Acquisition Without Dynamic Exclusion"

^3^Present Address: Clinical Research Center for Reproduction and Genetics in Hunan Province, Reproductive and Genetic Hospital of CITIC-XIANGYA, Changsha, Hunan, China

^4^Present Address: Molecular Medicine Program, The Hospital for Sick Children, Toronto, Ontario, Canada

*Corresponding Author.

**Table of Contents**

- **Cell line generation, culture, transfection, and collection**
- **Anti-FLAG AP of AP-MS samples**
- **Streptavidin AP of BioID samples**
- **MS data analysis (DDA)**

#### MS data analysis (DIA) and MSPLIT analysis settings

#### Interaction data analysis and visualization using ProHits-viz

- **Reference**
- **Table S1** Identification of *E. coli* proteins with different MS parameters.
- **Table S2** Gene ontology (GO) enrichment analysis of KRAS_G12V and KRAS_Q61H interactors.
- **Figure S1** The number of PSMs, peptides, and proteins identified in EIF4A2 and MEPCE AP-MS samples analyzed with the different methods.
- **Figure S2** Spectral count distributions of all high-confidence interactors for EIF4A2 and MEPCE.
- **Figure S3** Correlations between the log2-transformed spectral counts of all high-confidence interactors identified for EIF4A2 and MEPCE.
- **Figure S4** The number of PSMs, peptides, and proteins identified in BioID samples of mutated KRAS after analysis by the different methods.
- **Figure S5** Spectral count distributions of all high-confidence interactors for KRAS G12V and KRAS Q61H.
- **Figure S6** Correlations between the log2-transformed spectral counts of all high-confidence interactors identified for KRAS G12V and KRAS Q61H.

**Cell line generation, culture, transfection, and collection**

Stable cell lines for AP-MS (N-terminally 3×FLAG-tagged eukaryotic translation initiation factor 4A2 (EIF4A2) and methylphosphate capping enzyme (MEPCE)) and BioID (two N-terminally BirA*-FLAG-tagged activated forms (G12V, Q61H) of KRAS proto-oncogene, GTPase (KRAS)) were generated in HEK293 Flp-In T-REx cell (Invitrogen) pools as previously described^1^.

For both anti-FLAG AP-MS and BioID, one 15 cm plate was used per biological replicate. When cells reached 80% confluency in selection medium, they were induced with 1 µg/mL tetracycline for 24 h (for BioID, 50 µM biotin was added with the tetracycline). Cells were collected by removing the medium, washing them with phosphate-buffered saline (PBS), adding 1 mL PBS, and scraping the cells from the plate using a silicon cake spatula. For anti-FLAG AP-MS, ice-cold 1× PBS was used, and samples were kept on ice. Cells were pelleted at 500 × *g* for 5 min. The PBS was aspirated and the pellets were stored at -80°C.

**Anti-FLAG AP of AP-MS samples**

Tubes were transferred to ice. The cell pellets were weighed and resuspended at a 1:4 ratio (w/v) in lysis buffer (50 mM HEPES-KOH pH 8.0, 100 mM KCl, 2 mM ethylenediaminetetraacetic acid, 0.1% NP-40, and 10% glycerol, freshly supplemented with 1 mM phenylmethylsulfonyl fluoride, 1 mM dithiothreitol, and 1× protease inhibitor cocktail (Sigma-Aldrich)), then lysed by freeze-thaw by incubating the samples on dry ice for 5 min, then transferring them to a 37°C water bath until a small amount of ice remained. Samples were then centrifuged at 16,000 × *g* for 20 min at 4°C, and each clarified lysate was combined with 25 µL of a 50% slurry of anti-FLAG M2 magnetic beads (M8823, Sigma-Aldrich; pre-washed four times with 1 mL lysis buffer). Samples were gently agitated at 4°C for 2–3 h on a nutator. Beads were then washed twice with 1 mL lysis buffer and once with 1 mL rinsing buffer (20 mM Tris-HCl pH 8.0 and 2 mM CaCl_2_). After the final wash, the beads were centrifuged at 500 × *g* for 2 min and any remaining liquid was aspirated. Trypsin (750 ng in 20 mM Tris-HCl pH 8.0) was added to the beads, which were rotated at 37°C overnight. The following day, an additional 250 ng of trypsin (in 20 mM Tris-HCl pH 8.0) was added, and the sample was incubated at 37°C for 4 h. After the second trypsin digestion, formic acid was added to each sample to a final concentration of 5%. Peptides from five plates were combined and analyzed by noDE-DDA, DE-DDA, and SWATH acquisition methods in triplicate.

**Streptavidin AP of BioID samples**

Tubes were transferred to ice and the cell pellets were weighed then lysed by freeze-thaw as for AP-MS, but in 50 mM Tris pH7.5, 150 mM NaCl, 0.4% sodium dodecyl sulfate (SDS), 1% NP-40, 4.5 mM MgCl_2_ and 1 mM ethylene glycol tetraacetic acid, freshly supplemented with 1× protease inhibitor cocktail (Sigma-Aldrich) and 250 unit/mL TurboNuclease (Biovision). Samples were gently agitated on a nutator at 4°C for 30 min, then centrifuged at 16,000 × *g* for 20 min at 4°C. After centrifugation, each clarified lysate was combined with 25 µL of a 60% slurry of streptavidin Sepharose beads (17-5113-01, GE Healthcare; pre-washed four times in 1 mL in lysis buffer) and samples were gently agitated on a nutator at 4°C overnight. The next day, samples were centrifuged at 500 × *g* for 2 min, the supernatants were aspirated, and the beads were resuspended in 500 µL lysis buffer and transferred to fresh 1.5 mL Eppendorf tubes. The beads were washed once with 500 µL 2% SDS wash buffer (2% SDS, 50 mM Tris pH 7.5), twice with 500 µL lysis buffer, and three times with 50 mM ammonium bicarbonate pH 8.0 (ABC).

For protein digestion, trypsin (T6567, Sigma-Aldrich; 1 µg in 100 µL of ABC) was added and the beads were rotated for 4 h at 37°C. Then, an additional 1 µg of trypsin in 2 µL ABC was added and the beads were rotated overnight at 37°C. The tubes were centrifuged at 500 × *g* for 2 min and the supernatants were transferred to fresh 1.5 mL Eppendorf tubes. The beads were washed with 100 µL of high-performance liquid chromatography-grade H_2_O, and the supernatants were transferred to the same Eppendorf tubes to pool the peptides. Formic acid (50 µL of 10%) was added, for a final concentration of 2%. The pooled supernatants were centrifuged at 16,000 × *g* for 5 min to pellet the remaining beads, and 230 µL of each pooled supernatant was transferred to a new 1.5 mL Eppendorf tube and dried using a vacuum concentrator. The samples were then stored at -80°C until MS analysis, when the peptides were resuspended in 10 µL of 5% formic acid. For each MS analysis, 2.5 µL of the sample was used.

#### MS data analysis (DDA)

Mass spectrometry data generated were stored, searched, and analyzed using the ProHits laboratory information management system platform^2^. Within ProHits, WIFF files were converted to an MGF format using the WIFF2MGF converter and to a mzML format using ProteoWizard (V3.0.10702) and the AB SCIEX MS Data Converter (V1.3 beta). The data were then searched using Mascot (V2.3.02) and Comet (V2016.01 rev.2). The spectra were searched against the human and adenovirus sequences in the RefSeq database (version 57, January 30^th^, 2013) acquired from NCBI, supplemented with “common contaminants” from the Max Planck Institute (<http://lotus1.gwdg.de/mpg/mmbc/maxquant_input.nsf/7994124a4298328fc125748d0048fee2/$FILE/contaminants.fasta>) and the Global Proteome Machine (GPM; <https://www.thegpm.org/crap/>), forward and reverse sequences (labeled “gi|9999” or “DECOY”), sequence tags (BirA, GST26, mCherry, and green fluorescent protein (GFP)) and streptavidin, for a total of 72,481 entries. Database parameters were set to search for tryptic cleavages, allowing up to two missed cleavage sites per peptide with a mass tolerance of 35 ppm for precursors with charges of 2+ to 4+ and a tolerance of 0.15 amu for fragment ions. Deamidated asparagine and glutamine and oxidized methionine were selected as variable modifications. Results from each search engine were analyzed through the Trans-Proteomic Pipeline (v.4.7 POLAR VORTEX rev 1) via the iProphet pipeline^3^.

#### MS data analysis (DIA) and MSPLIT analysis settings

DIA data were analyzed using MSPLIT-DIA v1.0^4^. This approach was selected based on its high sensitivity for the detection of peptides/proteins present in the samples and direct benchmarking for protein-protein interaction detection. To generate a sample-specific spectral library, peptide spectral matches (PSMs) from matched DDA runs were pooled by retaining only the spectrum with the lowest MS-GFDB (Beta v1.0072, 6/30/2014)^5^ probability for each unique [peptide sequence](https://www.sciencedirect.com/topics/biochemistry-genetics-and-molecular-biology/peptide-sequence) and precursor charge state, and a peptide-level false discovery rate (FDR) of 1% was enforced using a target decoy approach. For AP-MS samples, DDA files from GFP, MEPCE, and EIF4A2 samples acquired using DE-DDA methods were used to generate the spectral library. For BioID samples, DDA files from all KRAS variants and control samples acquired by DE-DDA methods were used to generate the spectral library. The MS-GFDB parameters were set to search for tryptic cleavages, allowing no missed cleavage sites and 1 C^13^ atom per peptide with a mass tolerance of 50 ppm for precursors with charges of 2+ to 4+ and a tolerance of ± 50 ppm for fragment ions. Peptide length was limited to 8–30 amino acids. Oxidized methionine was selected as a variable modification. The spectra were searched against the NCBI RefSeq database (version 57, January 30th, 2013), containing 36,241 human sequences supplemented with adenovirus sequences and “common contaminants” as above. The spectral library was then used for protein identification by MSPLIT-DIA as previously described^4^, with peptides identified by MSPLIT-DIA that passed a 1% FDR subsequently mapped to genes using ProHits 4.0^6^. The MSPLIT-DIA search parameters were a parent mass tolerance of ± 25 Da and a fragment mass tolerance of ± 50 ppm. When retention time was available within the spectral library, a threshold of ± 5 min was applied to spectral matching as previously described^4^.

#### Interaction data analysis and visualization using ProHits-viz

Significance analysis of interactome (SAINT) express version 3.6.1 was used to score the enrichment of proteins in samples compared to negative controls using default parameters^7^. Two biological replicates of BirA*-FLAG-GFP, BirA*-FLAG-empty, and 3×FLAG-empty samples were used as negative controls for BioID experiments. Three replicates of 3×FLAG-GFP were used as negative controls for FLAG-AP experiments. Proteins with a Bayesian (B)FDR ≤1% and at least two unique peptides were considered true positive interactors. The entire network, scored with SAINTexpress, was used to generate input files for ProHits-viz^8^ for dot plot generation. To be included in a dot plot, a prey had to pass the 1% BFDR threshold with at least one bait. Once included, the prey’s quantitative values were recovered across all baits and methods, regardless of the BFDR. Unless otherwise indicated, the default options were selected, including ‘‘control subtraction’’ and hierarchical clustering using the Canberra distance metric and the Ward clustering type. Average spectral count values were capped at the maximum indicated in each figure, and unless otherwise indicated, no minimum spectral counts were required and no additional normalization steps were performed. When the entire network is not displayed in full, the rules governing the selection of the proximity interactions displayed are listed in the figure legends.

**Table S1.** Identification of *E. coli* proteins with different MS parameters (*n*=2)

| **Method** | **Collision energy** | **Q1 resolution** | **MS/MS acquisition mode** | **Top MS/MS candidate number** | **Precursor intensity threshold** | **MS/MS accumulation time** | **Proteins(#)** | **Peptides(#)** | **PSMs (#)** | **Sequence coverage** |
| --- | --- | --- | --- | --- | --- | --- | --- | --- | --- | --- |
| DE-DDA | Rolling | Unit | High sensitivity | 10 | 300 cps | 100 | 995 | 5086 | 10154 | 23.95% |
| noDE-DDA | Rolling | Unit | High sensitivity | 100 | 100 cps | 30 | 1108 | 6309 | 31773 | 29.00% |
| noDE-DDA fixed CE | Fixed | Unit | High sensitivity | 100 | 100 cps | 30 | 1063 | 6057 | 31302 | 28.25% |
| noDE-DDA low resolution Q1 | Rolling | Low | High sensitivity | 100 | 100 cps | 30 | 981 | 5483 | 28242 | 27.30% |
| noDE-DDA high resolution MS/MS | Rolling | Unit | High resolution | 100 | 100 cps | 30 | 910 | 5085 | 24852 | 25.25% |
| noDE-DDA top 75 MS/MS | Rolling | Unit | High sensitivity | 75 | 100 cps | 30 | 1088 | 6285 | 24360 | 29.55% |
| noDE-DDA 300cps | Rolling | Unit | High sensitivity | 100 | 300 cps | 30 | 1058 | 5971 | 30137 | 27.55% |
| noDE-DDA 45ms accumulation time | Rolling | Unit | High sensitivity | 100 | 100 cps | 45 | 1085 | 6046 | 22535 | 28.05% |
| noDE-DDA 60ms accumulation time | Rolling | Unit | High sensitivity | 100 | 100 cps | 60 | 1049 | 5724 | 17177 | 27.20% |

**Table S2.** Gene ontology (GO) enrichment analysis of KRAS_G12V and KRAS_Q61H interactors that passed SAINTexpress filtering (BFDR ≤ 1%). BP, biological process; MF, molecular function; CC, cellular compartment.

| **Term (Source)** | **DE-DDA** | | | | **MSPLIT-DIA** | | | | **noDE-DDA** | | | |
| --- | --- | --- | --- | --- | --- | --- | --- | --- | --- | --- | --- | --- |
|  | KRAS G12V | | KRAS Q61H | | KRAS G12V | | KRAS Q61H | | KRAS G12V | | KRAS Q61H | |
|  | *p*-value | Ranking | *p*-value | Ranking | *p*-value | Ranking | *p*-value | Ranking | *p*-value | Ranking | *p*-value | Ranking |
| Signal transduction (BP) | 6.72E-09 | 75 | 3.13E-07 | 87 | 7.61E-08 | 108 | 2.75E-06 | 134 | 9.75E-12 | 69 | 1.39E-12 | 67 |
| Ras protein signal transduction (BP) | 3.26E-06 | 138 | 2.14E-04 | 172 | 3.78E-05 | 190 | 5.46E-03 | 289 | 1.30E-07 | 136 | 4.10E-09 | 113 |
| Regulation of molecular function (BP) | 4.04E-05 | 174 | 1.25E-04 | 165 | 1.18E-06 | 135 | 2.96E-05 | 171 | 3.96E-09 | 105 | 4.04E-14 | 60 |
| Regulation of GTPase activity (BP) | 2.21E-11 | 50 | 2.11E-07 | 77 | 1.68E-09 | 76 | 1.58E-06 | 122 | 5.71E-18 | 38 | 8.13E-22 | 18 |
| Protein binding (MF) | 6.41E-13 | 3 | 1.87E-14 | 3 | 1.05E-12 | 3 | 2.95E-14 | 3 | 8.17E-17 | 4 | 2.15E-19 | 3 |
| Ras GTPase binding (MF) | 2.45E-07 | 13 | 1.34E-07 | 15 | 3.71E-08 | 15 | 1.31E-05 | 30 | 1.75E-25 | 6 | 1.10E-15 | 7 |
| Plasma Membrane (CC) | 4.69E-58 | 2 | 3.78E-51 | 2 | 4.36E-58 | 2 | 4.92E-55 | 2 | 1.45E-70 | 2 | 8.98E-64 | 2 |
| # of proteins | 329 | | 304 | | 323 | | 292 | | 452 | | 419 | |

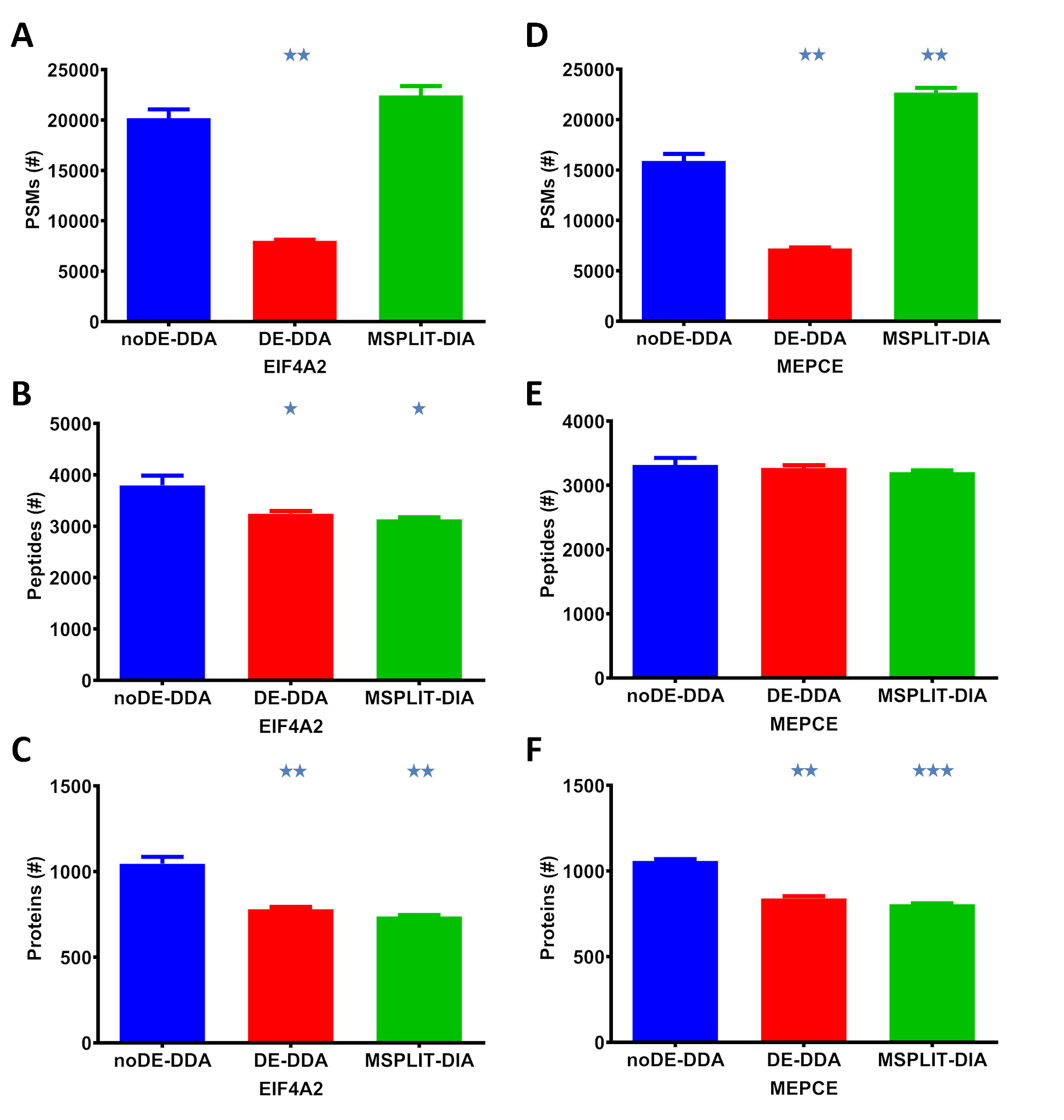

**Figure S1.** The number of PSMs (A and D), peptides (B and E), and proteins (C and F) identified in EIF4A2 and MEPCE AP-MS samples analyzed with the different methods (*n*=3, average). **p* < 0.05, ***p* < 0.01, and ****p* < 0.001, by two-tailed unpaired Student’s *t*-test.

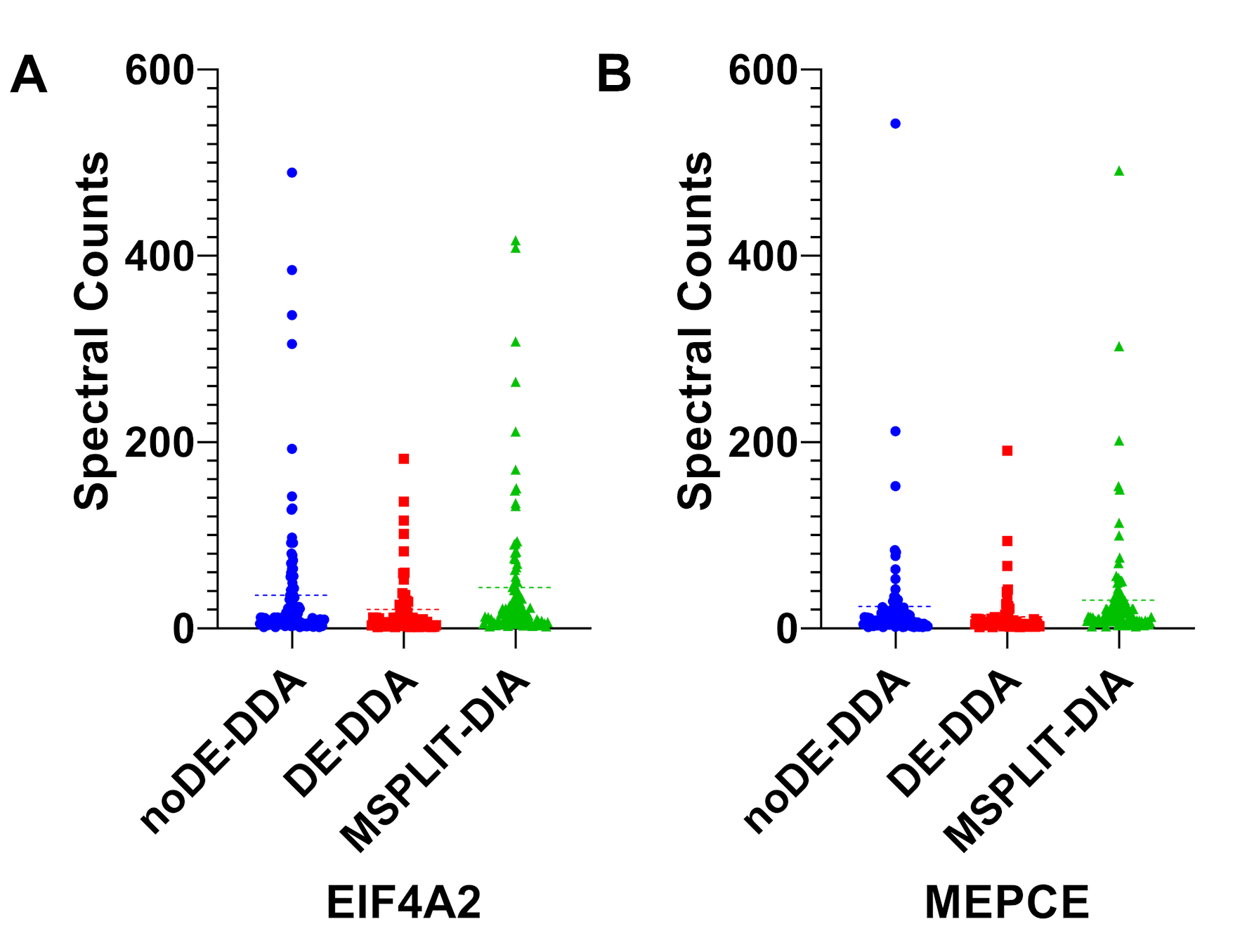

**Figure S2.** Spectral count distributions of all high-confidence interactors for EIF4A2 (A) and MEPCE (B) identified by the different methods after filtering by SAINTexpress (BFDR ≤ 1%).

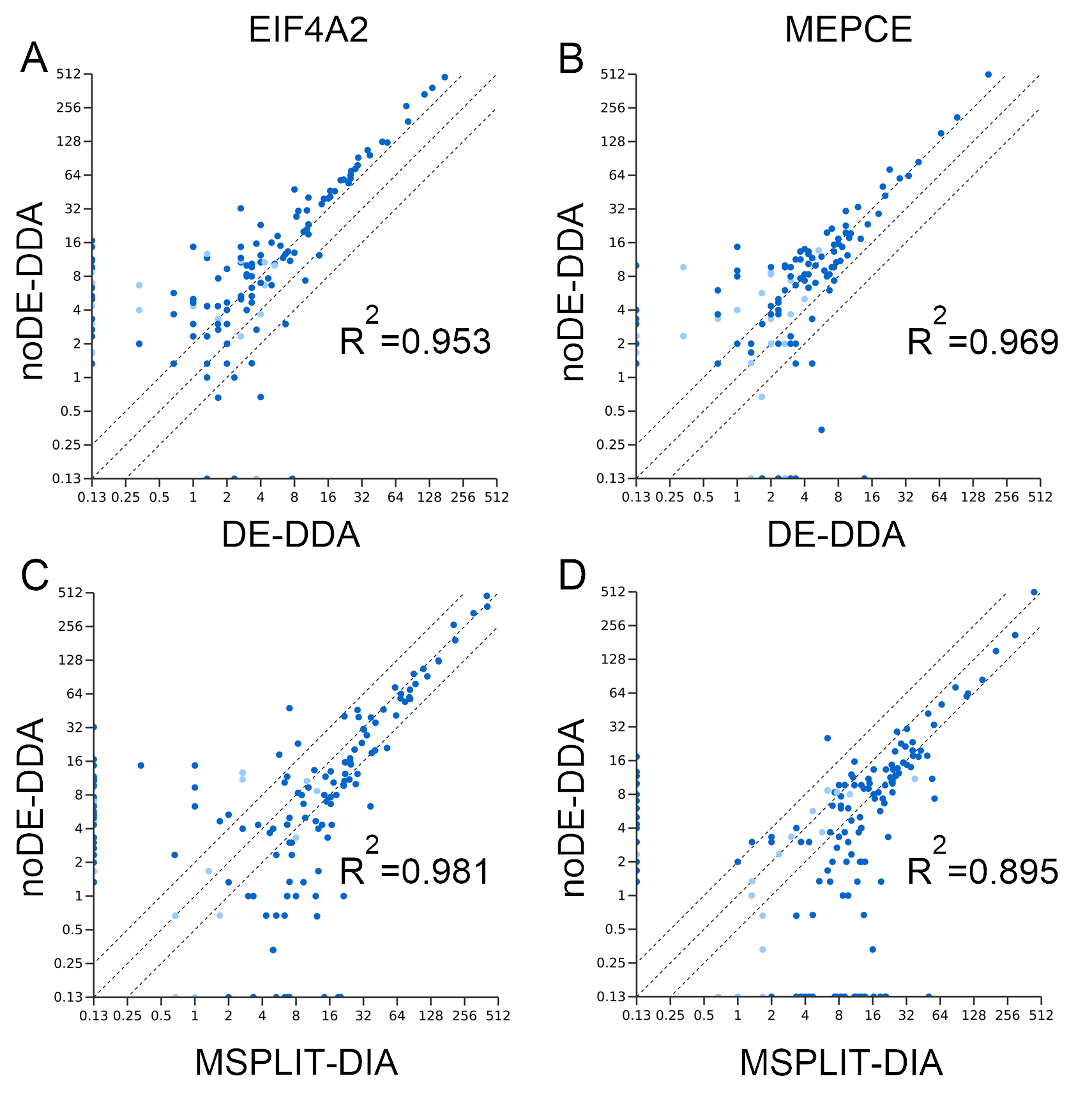

**Figure S3.** Correlations between the log2-transformed spectral counts of all high-confidence interactors identified for EIF4A2 (A and C) and MEPCE (B and D) by the different methods after filtering by SAINTexpress (BFDR ≤ 1%).

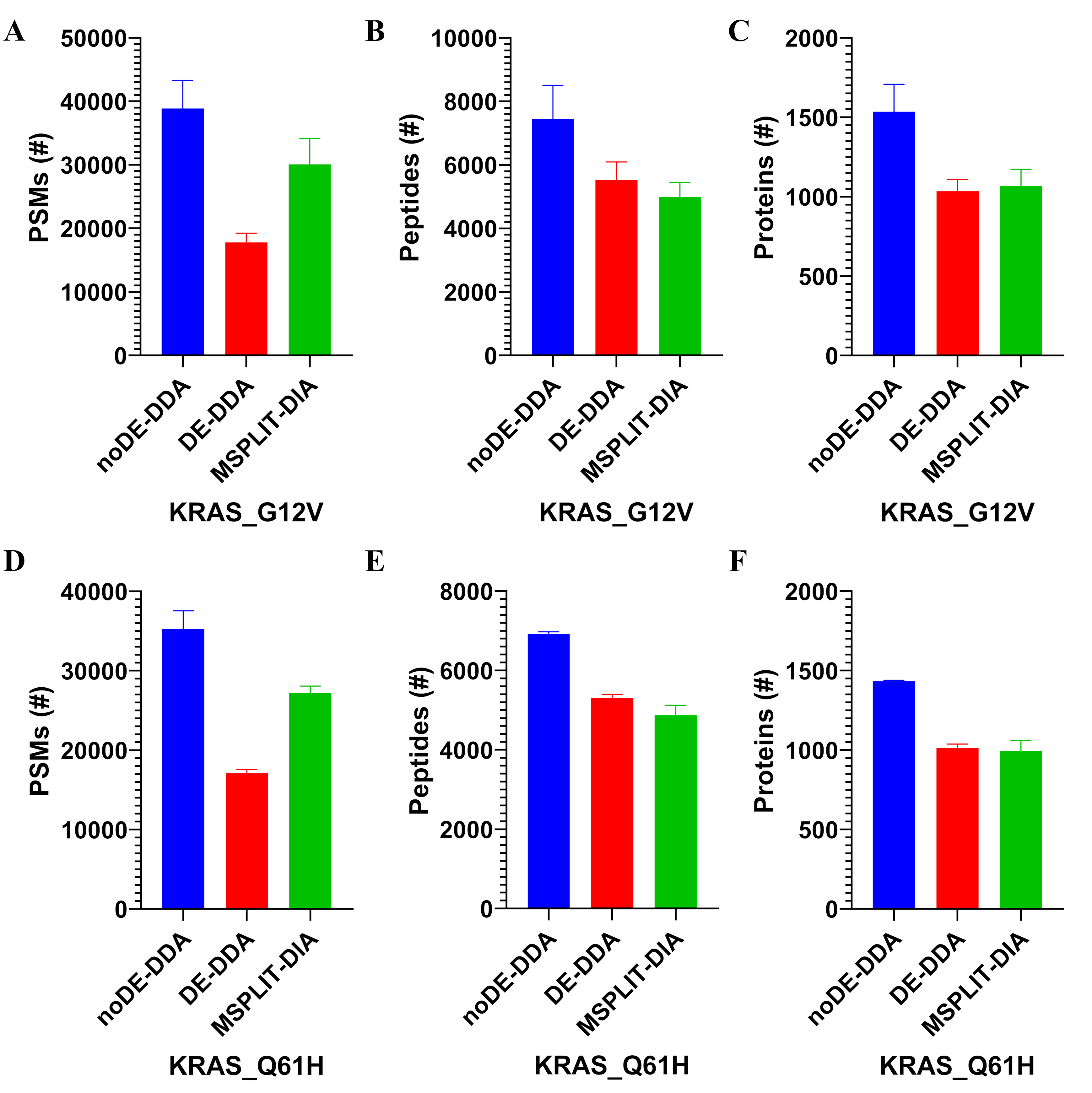

**Figure S4.** The number of PSMs (A and D), peptides (B and E), and proteins (C and F) identified in BioID samples of mutated KRAS after analysis by the different methods (two biological replicates).

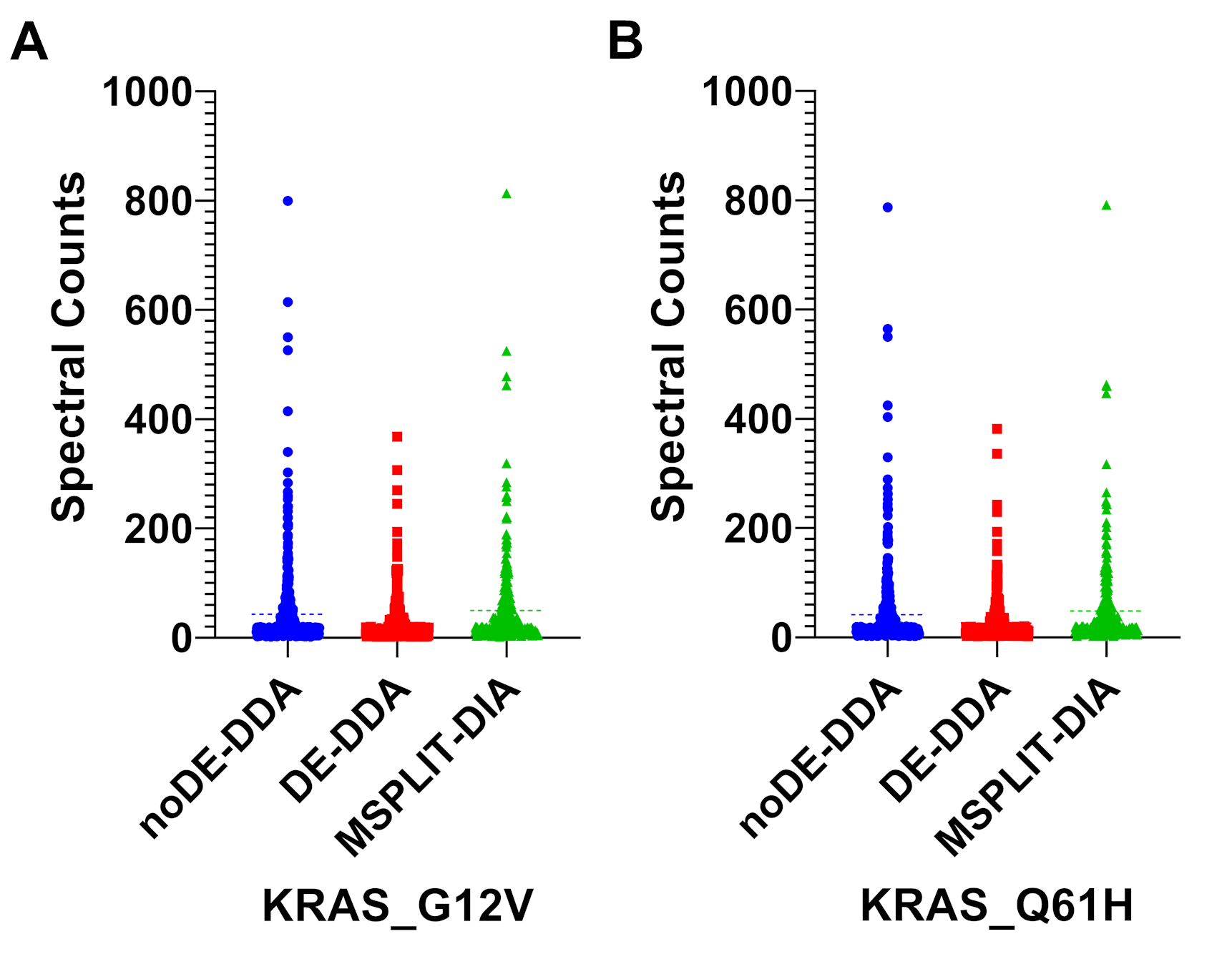

**Figure S5.** Spectral count distributions of all high-confidence interactors for KRAS G12V (A) and KRAS Q61H (B) identified by the different methods after filtering by SAINTexpress (BFDR ≤ 1%).

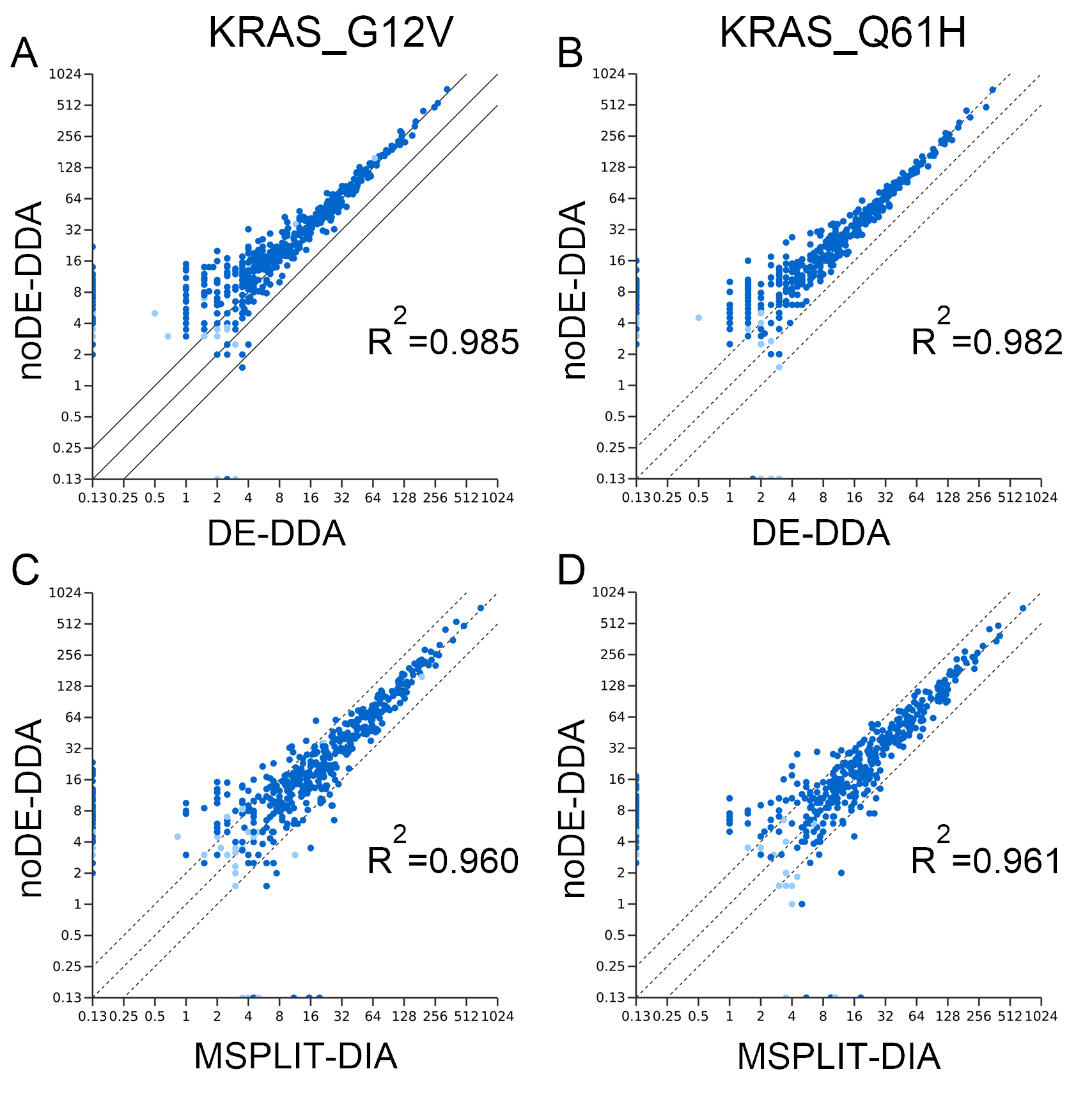

**Figure S6.** Correlations between the log2-transformed spectral counts of all high-confidence interactors identified for KRAS G12V (A and C) and KRAS Q61H (B and D) by the different methods after filtering by SAINTexpress (BFDR ≤ 1%).
